# Fraudulent studies are undermining the reliability of systematic reviews – a study of the prevalence of problematic images in preclinical studies of depression

**DOI:** 10.1101/2024.02.13.580196

**Authors:** Jenny P. Berrío, Otto Kalliokoski

## Abstract

Systematic reviews are considered by many to constitute the highest level of scientific evidence. A caveat is that the methods used in a systematic review – combining information from multiple studies – are predicated on all of the reports being truthful. Currently, we do not know how frequent fraudulent studies are in systematic reviews, or how they affect the resulting evidence base. For a systematic review of preclinical studies of depression, we found that potentially fraudulent studies were not only common but also that they biased the findings of the review. In a sample of 1,035 studies, we found that 19 % of peer-reviewed reports displayed data in the form of problematic images. In a majority of the cases, images had been altered or recycled in a way that makes us suspect foul play. Making things worse, these studies reported larger effect sizes, on average, than did studies where we did not identify problems. Counter to commonly held beliefs, reports with problematic images were not cited less or published in lower-impact journals, nor were their authors isolated to any specific geographic area. The sheer prevalence of problematic studies, and the fact that we could not find a simple pattern for identifying them, undermines the validity of systematic reviews within our research field. We suspect that this is symptomatic of a broader problem that needs immediate addressing.

## Introduction

Systematic reviews are considered by many to be the pinnacle of medical evidence^1,2^, if carried out properly. Systematic reviews of clinical studies inform best practices with respect to patient care; systematic reviews of preclinical studies are used to elucidate fundamental biological mechanisms, but also to inform decisions on which drug candidates to develop and which clinical trials may be worthwhile to carry out^3,4^. Numerous guidelines and best practices have been published to aid researchers in combining and synthesizing evidence in the best possible way^5^. There is however no broad consensus on how to deal with potentially fraudulent studies^6^. When should we exclude a study simply because we do not trust it?

Can we do so without introducing an element of bias? Most guidelines trying to gauge the reliability of findings in a peer-reviewed report assume that the authors are acting in good faith. A study took place, and it was conducted as written. But, what do we do when we suspect that this is not the case?

While conducting a systematic review of preclinical studies of depression, we found publications displaying research data using problematic images. Many of these images showed evidence suggestive of fabrication or falsification. In the present report, we have systematically assessed and documented the types and frequencies of issues we encountered in our screening process. Based on our findings, it seems fraudulent studies are more frequent than is suggested by many previous estimates^7^. They are also potentially warping the conclusions of our systematic reviews. Moreover, traditional methods for avoiding low quality studies are ruefully inadequate in identifying problematic papers. Focusing on the renown of the journal publishing the paper, or seeing whether the findings were cited by other researchers, does not seem to reduce the chances that a paper presents with problematic images. Checklists and protocols for gauging the quality of evidence in a systematic fashion^8,9^ also do not single out these problematic reports. If we are to retain the integrity of meta-analytical investigations in general, and systematic reviews in particular, we desperately need to develop better methods for identifying and dealing with potentially fraudulent reports in peer-reviewed journals.

Our systematic review concerned unpredictable chronic stress, and its use in modelling depression in laboratory rats^10^. The theory is that a rat exposed to constantly-shifting-but-invariably-stressful conditions on a daily basis will develop something akin to a major depression^11^. The method has been employed across thousands of experiments^12^, on what must be more than a million rats by now. Yet, we are not certain to what extent it is a good method for studying depressions^13^, if at all. The field is consequently ripe for meta-analytical investigation. We were interested in evaluating the efficacy of studying the effects of this stress protocol using the sucrose preference test – a test that measures the ability to feel joy by studying how rats, when given the chance, will binge on a sugary solution^14^. There are a number of different approaches within this model so, to work with a uniform set of studies, we further constrained our investigation. We elected to combine data from only experiments that applied the stress paradigm for at least two weeks, and where the rats were not excessively fasted (< 6 h) before the sucrose preference test.

Peer-reviewed publications were sourced through journal databases. After an initial screening step, we were faced with the task of skimming a large number of publications to evaluate whether they fit the scope of our investigation. At this stage, we noticed inconsistencies and duplications in images presenting data. Not knowing how to conscientiously deal with the associated publications, we chose to document these issues separately.

To be able to speak cogently about the issues we found, we have used a classification system developed by Dr. Elisabeth Bik and collaborators^15^. The system distinguishes between three levels of issues – those that could easily have arisen by accident in preparing the paper; those where the issues could have arisen by accident, but not easily; and issues that stem from images having been intentionally manipulated (**Figure 1**). We will refrain from speculating why images had been altered, save for noting that we analyzed only images presenting research data – changing them in any way is tampering with this data and, by its very definition, constitutes research misconduct^16,17^.

**Figure 1.**
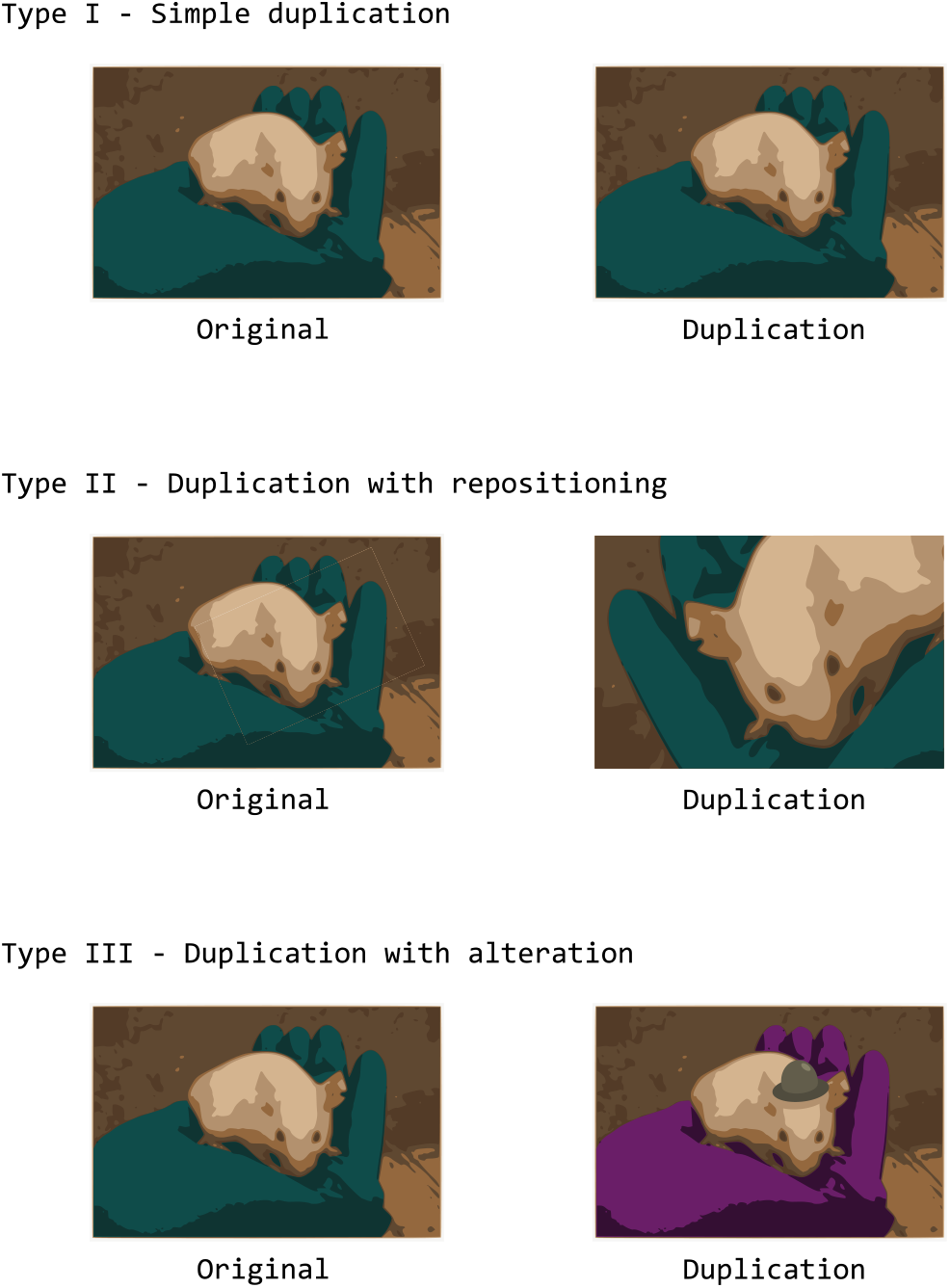
Classifying image issues using the “Bik scale.” Using examples, we can define three levels of problematic image duplications^15^. Simple duplications (Type I) are when an image occurs twice in the same report, but ostensibly portrays two different things. If only parts of an image is duplicated and presented as something else, we consider this a duplication with repositioning (Type II). In the latter scenario, we often see that the duplicated image has been rotated and/or mirrored, almost as if to obscure the duplication. This is demonstrated here, where we have highlighted the re-used image segment, which has been both rotated and mirrored. When image-editing software has been used to remove or add features to an image, or to fuse elements from multiple images, we consider this a duplication with alteration (Type III). Whereas none of these issues should occur in a report, only the last type is a clear expression of deception (falsification/fabrication). We have used a simple, striking, image for demonstration purposes. Note that the image duplications we discuss below are much harder to identify.

In addition to presenting frequencies and types of issues, we present how studies with problematic images affected our systematic review. We also conducted a simple bibliometric analysis to address questions that often arise when discussing potentially fraudulent studies.

## Results and discussion

### Overview

In our investigation, we scanned 1,035 peer-reviewed publications, but only 588 papers presented research data in image form (excluding line drawings like plots, graphs, infographics, and schematics). We flagged 112 papers with problematic images, suggesting that one in five papers within the research field (19 %) suffer from these issues (**Figure 2**). Alarmingly, the problematic images we found were heavily skewed toward types II (49 studies) and III (33 studies), suggesting that a large number – we suspect the majority – of the issues are evidence of manipulation of primary data. The simple duplications – Type I – that could somewhat easily happen in the final stages of preparing a manuscript were, by contrast, rare (13 studies). In addition, we found some issues that could not be slotted into the three categories (Other issues: 17 studies). These ranged from benign mistakes where images had likely been switched around, producing mismatched descriptions, to papers where the images bore the fingerprints of a suspected “paper mill” (individuals fabricating research reports for payment^18^).

**Figure 2.**
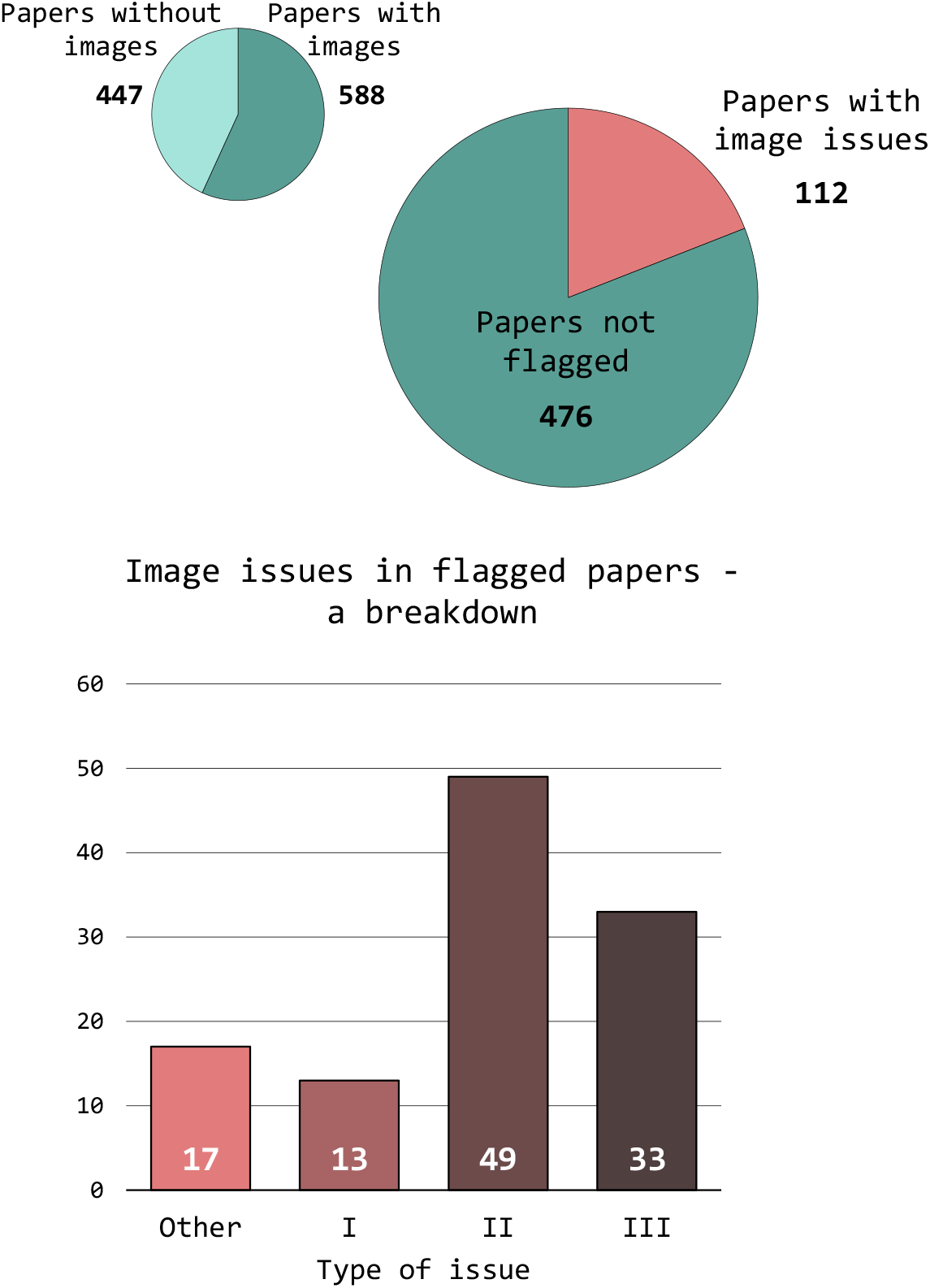
Breakdown of 1,035 papers that were screened for a systematic review of preclinical depression studies. Slightly more than half (57 %) of the papers presented one or more types of data in the form of images (not including line drawings like plots or schematics). Of these, 19 % contained at least one problematic image. Alarmingly, the types of problems that we identified were heavily skewed toward issues suggestive of willful manipulation of the portrayed research data (Types II and III). For a summary of the types of issues, refer to **Figure 1**.

The image issues were found across numerous representations of research data – from path diagrams ostensibly displaying how an animal traversed a particular behavior testing apparatus to photographed gels of PCR products. Most commonly, duplications were found in western blot images (54 studies). Issues with micrographs were also frequent, with histology images (28 studies), immunohistochemistry images (18 studies), and immunofluorescence images (13 studies) being particularly numerous. It is also worth noting that some studies (11 studies) contained problems spanning multiple image types.

Although only a third (29 %) of the studies featured clear alterations (Type III issues), many of the problematic reports in the other categories were suspicious to us. In many cases, it would be difficult to construct a reasonable-sounding narrative that might explain the issues we found as the results of carelessness. The numerous Type I duplications in some reports would be quite difficult to simply explain away as a very unlucky individual (with equally distracted co-authors) being absentminded – repeatedly (one report featured 14 pairs of duplicated images) – when compiling the report. Similarly, some of the Type II duplications were so involved, with re-scaling, rotation, mirroring and contrast adjustments of the images, that it was difficult to think of them as anything other than deliberate strategies employed to conceal foul play. Consequently, it is our professional assessment that a majority of the issues we found were indicative of misconduct.

### Effect on meta-analyses

Of the 1,035 papers we screened, 132 met all of our requirements for being included in our systematic review. Ten of these were flagged with image issues; six of them with Type II (3) or Type III (3) issues. For our meta-analyses, we were interested in how the sucrose intake of rats exposed to the chronic stress paradigm compared to unstressed controls. This was rarely the central comparison in the papers we were synthesizing. The sucrose preference test is often used simply to verify that the stress-induced depression model is working, frequently as part of a panel of tests. Yet, perhaps unsurprisingly, all but one of the papers flagged with image issues found a statistically significant difference between stressed rats and controls. Overall, the flagged papers presented with effect sizes that were, on average, higher than that of papers that we had not flagged (Average difference: 0.81 standard deviations; Cochran’s Q test used for sub-group analysis: Q_1_ = 4.57, p = 0.03), but not so much so that they would look out of place (**Figure 3**). In many ways, the studies were insidiously average. Neither our checklists for quality of reporting nor risk of bias set them apart from the other studies. Eight studies that were flagged with image issues were scored for their quality of reporting, using a checklist based on the ARRIVE guidelines^19^. On average, these studies scored a 5.8 out of ten (range: 3-9), where a higher number indicates a better quality of reporting. The overall average for all 100 studies that were scored for their quality of reporting was 5.1. Similarly, all ten of the studies flagged with image issues were graded using a common tool for assessing the risk of bias stemming from the design of the study^8^. On average, the studies scored a 3.2 out of eleven (range: 2-5), which suggests that there was a high risk of bias – a higher score indicates that more efforts have been taken to mitigate sources of bias. Unfortunately, this is simply symptomatic for the research field – the average score for all 132 studies included in the review was 2.9.

**Figure 3.**
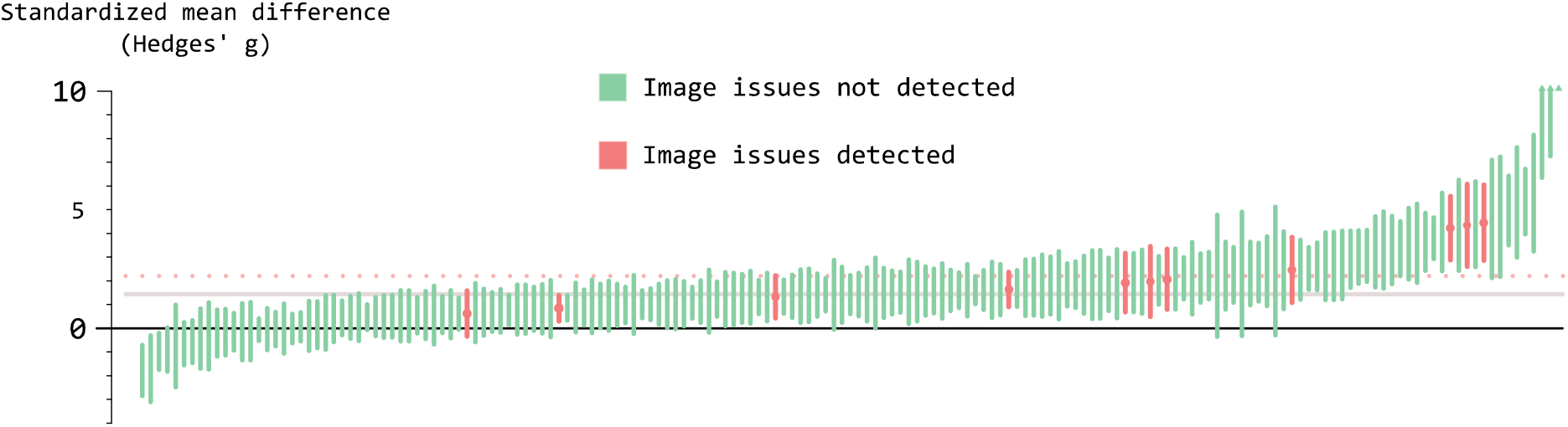
Effect sizes in the studies included in the systematic review. Each vertical line represents the 95 % confidence interval of the standardized mean difference found between the stressed group and the control group in an experiment. The greater the value on the Y-axis, the greater the difference between stressed rats and control rats in the sucrose preference test (i.e. the less interest the stressed rats showed the sugary solution compared to controls). We have highlighted eleven experiments from ten papers that were flagged with image issues. The gray horizontal line represents the average effect across all studies. The dotted horizontal line is the average of the flagged papers. Three studies presented average effect sizes greater than 10 standard deviations; they are consequently not shown in full in the figure. None of the three was flagged with image issues, although it can be argued that the effect sizes described in the papers are implausibly large.

At this point, it is worth noting that we are only discussing one type of problematic studies. There is no reason to believe that reports presenting data in image form are more prone to being problematic. If we consider fraudulent studies, for example, it is more difficult to manipulate images than it is to falsify tables, fabricate bar charts, or plagiarize text. We have not made any attempts in the present investigation to check statistics^20^ or tabulated numbers^21^ for internal consistency, searched for plagiarized snippets of text^22^, or employed any of a number of other methods^23^ proposed for detecting problematic reports. Our methods would not even have been able to find competently manipulated images. A skilled Photoshop user could easily fool us, and the image analysis software we used. As high as our numbers are, they are still a low estimate for potentially fraudulent papers.

### Bibliometrics

A common misconception is that fraudulent studies are nigh on exclusively found in so-called predatory journals – journals lacking proper peer-review and editorial quality checks^24^. These have been described as “reservoirs of author misconduct^25^” by Jeffrey Beall (of Beall’s List fame). But were our reports with problematic images more likely to be found in journals of ill-repute? Our bibliometric analysis did not reveal any differences between the impact factor of journals publishing papers with problematic images (where we suspect a majority may be fraudulent) and a random sample of papers where we detected no image issues (**Figure 4**). We are the first to admit that impact factors are a poor metric of a journal’s quality (as others have in the past^26,27^). Nevertheless, if the fraudulent studies were principally published in journals with poor renown, like predatory journals, we would have expected to find a difference. Here, our data contradicts the findings of Bik *et al*., who found a trend with reports containing problematic images being more frequent in journals with a lower impact factor^15^. The diverging results may be an effect of us focusing on a publications level, whereas the previous investigation chose to focus on a journal level.

**Figure 4.**
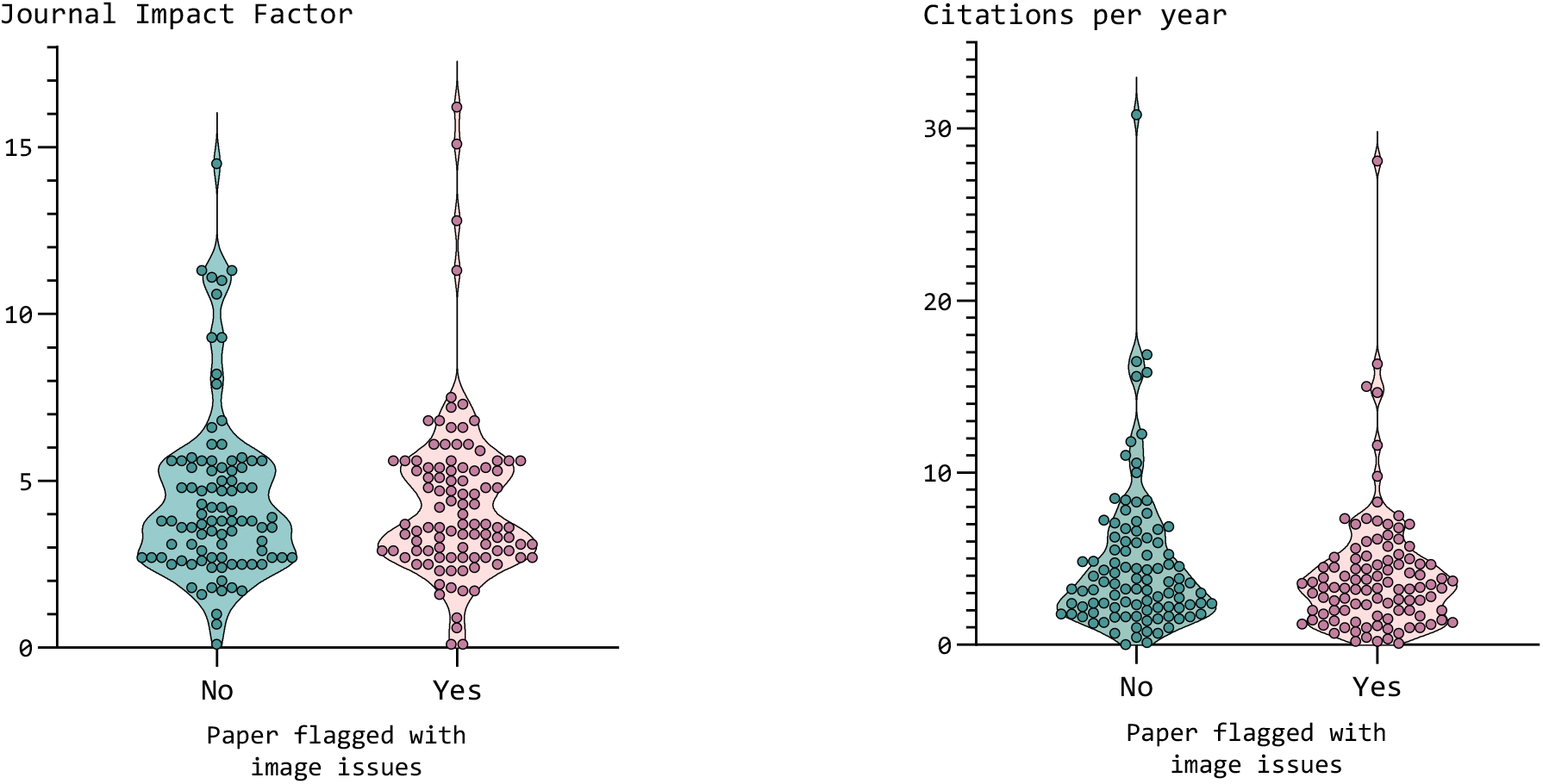
A simple bibliometric analysis suggests that papers with problematic images cannot be distinguished from papers using simple metrics. Neither the impact factor of the journals that publish these papers (median: 3.3), nor the yearly number of citations they accrue (median: 3.7), can be used to identify problematic and potentially fraudulent studies.

Another common misconception is that researchers know instinctively to avoid relying on fraudulent studies. When discussing these issues with colleagues, many claim that they could never have cited or relied on a potentially fraudulent report – they will insist that “they have a nose for these things”, or something to that effect. The analysis, again, does not support this. On average, the studies featuring problematic images were cited as often as were the studies without issues.

It should be noted that we could also find no difference in bibliometric trends between studies featuring different types of issues. The absence of any (statistically substantiated) difference, with respect to both journal impact factors and yearly citations, remained also when we, for example, restricted the analysis to only papers with Type III issues.

Clearly, we cannot avoid problematic studies simply by shunning rarely-cited publications (relying on “the wisdom of the crowd”) or low-impact journals wholesale. Another strategy for avoiding potentially fraudulent papers is completely ignoring publications coming from countries considered hotbeds for “paper mills.” China has been implicated to be such a country on multiple occasions^28-30^. Whereas our dataset is not large enough to allow for a thorough analysis of author affiliations, we can make a few observations.

Outside of the obvious issues of discrimination/outright racism^31^, there are additional reasons for why ignoring publications from *e*.*g*. China is a bad heuristic. Of the 1,035 papers we screened, 607 (58.6 %) had one or more authors with a Chinese affiliation. Ignoring all publications coming out of the world’s largest producer of scientific studies^32^ is simply not feasible. China has admittedly struggled with poorly aligned researcher incentives^33^ fostering a problematic publication culture in certain research environments^34^. This is also evident in our data where a study was more likely to have problematic images if it had one or more authors with a Chinese affiliation (risk ratio: 1.42). But, so were studies authored by researchers with a Danish affiliation. Of 24 studies containing analyzable images and with one or more Danish authors, seven were found to have problematic images (risk ratio: 1.98). The authors of the present report would not like to be judged based on a productive laboratory on the other side of the country producing a number of studies with problematic images. We are sure that this sentiment is one that is shared by many of our Chinese colleagues.

### Responses and reactions

The topic of fraudulent studies, and facing their prevalence, is uncomfortable and inconvenient. Using bibliometric information to judge a study’s veracity comes across as a desperate attempt at circumventing the problem without having to deal with it directly. Clinging to simple solutions that would retain the *status quo* is as understandable as it is misguided. It is also common to see discussions about fraudulent studies diverted with an appeal to more widespread issues. Questionable research practices, underpowered studies, and statistical illiteracy are more deserving of attention, we will invariably be told. Whereas we do not disagree with these being pressing issues – not in the least! – it does not diminish the need for a consensus on how to identify and deal with fraudulent studies. Our investigation found image issues in one-in-five peer-reviewed papers, the majority of which we believe to be, in parts or fully, doctored. No amount of teaching the proper uses of a t-test will address this.

We would suspect that the reason why most researchers would agree with the statement that research fraud is rare, all the while data seems to point to the contrary^35,36^, is that they have not encountered it personally^37^. “If I have never seen anyone commit fraud, how could it be common?” We believe that the missing puzzle piece is to understand that fraud is likely conducted by a very small number of people. In our investigation, we found that many of the flagged papers stemmed from the same laboratory, with recurring authors. This is consistent with previous investigations of research fraud where, once issues indicative of fraud are uncovered in a publication, it is often subsequently found that one or more of the authors are repeat-offenders^38,39^. Since authors willing to falsify their results are not subject to the cruel whims of chance but always guaranteed to obtain the desired results in their experiments, they can be very productive. Authors who are ready to make up their results without any experiments whatsoever, and those who employ “paper mills” to do this for them, can be even more productive. This is how a small number of people^40^ with flexible morals can pollute the scientific literature disproportionately. It is also a piece of information that offers a sliver of hope in the face of very bleak numbers. We can reinstate reliability in a given research field by identifying a relatively small number of culprits and retracting their work. This is conditional on a willingness to take action, however.

Since reporting 107 problematic studies on the post-publication peer review platform PubPeer (five papers could not be reported due to issues concerning the digital object identifiers (DOIs)), only a single publication has been retracted. The paper in question featured what we characterized in our notes as

“pervasive use of Photoshop across multiple images” and the paper was flagged as deeply problematic on PubPeer, before we had the time, by another anonymous user using the identity *Hoya camphorifolia* (anonymous PubPeer users are assigned a random user name from a taxonomic database). The paper is suspected to be the product of a particularly prolific paper mill^41^. The retraction was requested by the journal’s Editor-in-Chief, after they had unsuccessfully sought the (purported) authors for an explanation^42^. In the case of another twelve papers, errata/corrigenda have been issued where the images we had pointed out have been replaced with images without duplications. This includes eight cases where the original images had substantial issues (Types II and III). In most cases, no explanation to the original problems has been offered. Some authors (another 12 cases) have chosen to respond to the issues through PubPeer. In only one case has an author disagreed with the core issue, arguing that the images were, in fact, not duplicated. In this case, the authors claimed to possess high-resolution scans of the original western blots, proving that the images were only highly similar but not actually duplicated. These scans were not shared, whereby we consider the issue still to be unresolved. In five cases, the authors claim to have contacted the journal for correcting the errors in question, but without a corrigendum having been issued. Some of these claims are more than two years old at the time of writing, leading us to wonder whether the issues will ever be corrected. In only one case did the authors issue a correction^43^ providing a transparent, plausible, and collegial explanation to the issues (a Type I duplication) we had raised. We applaud the authors’ model behavior and wonder why this happened only once in 107 cases. Most authors and journals have chosen not to engage with the reported issues entirely. The studies originating from a Danish research group were reported to the Danish Board on Research Misconduct. The researchers were cleared on all five counts of suspected misconduct^44^. The investigation consisted of contacting the researchers and asking about the images, which they insisted were not duplicated. Impartial experts were not consulted, the papers remain uncorrected, and the ruling cannot be appealed. Outside of facing the problem of a deluge of potentially fraudulent studies, we are also facing a problem with an institutional unwillingness to deal with potentially fraudulent studies.

## Conclusions

Our screenings show that peer-reviewed reports with problematic images are common within the field of preclinical depression studies. We believe that a majority of these reports had been, in part or completely, fabricated or falsified. Moreover, in the context of our systematic review, they served to inflate our meta-analytical estimates. We could also not find a simple pattern that would allow for detecting, and potentially excluding, these problematic reports outside of carefully investigating their images. The consequences of our findings are concerning. Any preclinical systematic review and meta-analytical investigation carried out in this field will potentially be misled by fraudulent studies.

We may ask ourselves whether these results are specific to the field of preclinical depression studies. Without more data, we cannot give a certain answer, but there is reason to believe that our numbers can be extrapolated to other preclinical research fields. Firstly, the chronic unpredictable stress model is widely used across sub-fields. It is used to investigate fundamental neurobiological mechanisms, to study depression comorbidities, to test antidepressant drug candidates, and in many more contexts. The reports with problematic images do not seem to be isolated to any one of these sub-fields. The problem seems to affect preclinical neurobiology/neuropharmacology at large. Secondly, our estimate for potentially fraudulent papers aligns well with recent discoveries. There are few investigations that have employed similar approaches; however, the ones we know of^15,45-48^ have produced similar estimates for the prevalence of reports featuring problematic images (although we appear to have the dubious honor of having produced one of the highest estimate so far).

What then is the potential damage? Focusing on a small number of high-profile fraud cases may give the appearance that fraudulent studies are used to create sensational findings – major advances in cloned human stem cells^49^, discovery of novel superconductors^50^, finding the root cause of Alzheimer’s disease^51^, harnessing the power of social priming^52^, *etc*. In the context of systematic reviews of human trials, much focus has been on a small number of studies reporting extreme effects. Arguably for good reasons^53,54^. Yet, what we see in our investigation is that – at least within preclinical studies – most problematic studies (and among them, we speculate, the fraudulent) are mundane. They do not to make waves; they agree with the general consensus within the field. Sure, we found that reports with problematic images present higher-than-average effect sizes, but they were not the highest. It can be argued that these reports constitute just as a big a problem, since they cannot easily be detected. Even if the individual studies do not present hyperbolic findings, they muddy the waters, making truthful effects harder to gauge with any accuracy. In the context of preclinical depression studies, we already have to contend with evidence of publications bias^10^ and exaggerated effects^55^. Fraudulent studies going along with these inflated effects ensure that erroneous findings become difficult, if not impossible, to overturn. It can be shown that the “self-correcting” nature of science is – at best – torturously slow^56,57^. Problematic papers may serve to completely derail this process.

Simple shortcuts to determining the reliability of a report do not seem to exist. We might be tempted to rely on studies that have often been cited in the past. However, citations are more likely to reflect whether the results of a study are convenient – for example, by fitting a particular narrative or a theory – than being a metric of methodological quality, theoretical rigor, or overall veracity. We might set our hopes to the integrity of esteemed journals. But, again, we would probably be misled. Whereas journals with a higher impact factor appear to be less likely to publish a study with problematic images^15^, our investigation suggests that being published in a higher-impact journal is no guarantee of a paper’s integrity. At the end of the day, each study has to be judged on its own merits.

Currently, Cochrane argues in their guidelines that retracted studies should be excluded from systematic reviews^58^. We argue that this is insufficient. Journal retractions happen at a glacial speed, and author protests can sometimes be enough to prevent them wholesale^59,60^. The scientific community cannot afford to act as slowly as the industry that publishes our reports. Our ability to collaboratively discover the underlying causes of disease and efficacious treatments thereof in the field of preclinical medicine is undermined by falsified and fabricated evidence. We would argue that any report featuring inappropriate duplications of images – no matter the type of duplication – should be excluded from evidence synthesis in both meta-analyses and systematic reviews. It might seem inadvisable to exclude an important study from a systematic review merely because of a mix-up concerning its images; a case of throwing the baby out with the bathwater. But, we should remember that it is quite simple to correct a publication if a duplication stems from an honest mistake. It is trivial for authors to provide a transparent account of what has happened, how this affects the results, and to provide a corrected report. If we wish to preserve the systematic review’s status as the pinnacle of research evidence, drastic steps must be taken to combat the influx of potentially fraudulent studies. Putting the onus on the authors to correct their report if it is to be made a part of the evidence base of a research field does not seem an unreasonable approach to us. For the longest of time, we have been laboring under the assumption that dishonest reports are virtually non-existent. The unfortunate truth is that this is not a position we can maintain.

## Methods

### Screening

Peer-reviewed publications were sourced for our systematic review through EMBASE, PubMed and Web of Science. For details on search strings and initial screening steps, refer to the original publication^10^.

Importantly, papers not describing original experimental studies – *i*.*e*., reviews, meta-analyses, opinion papers, and meeting abstracts – were excluded in an initial screening step. Of the 1,035 items that were included in the full text screenings, we singled out studies that featured one or more photograph/micrograph/scan presenting research data. We also included reports with either flow cytometry plots or path diagrams from behavioral experiments, since there are previous examples of these having been doctored. Although this is also true for spectroscopic or chromatographic spectra from *e*.*g*. Raman spectroscopy^61^, we excluded these since we were not confident in our ability to separate similar images from identical images in low-resolution images of spectra. Furthermore, we did not consider example images that did not portray original data. Images of experimental setups and stylized images demonstrating a method can, obviously, be edited without this affecting the reliability of the associated report.

In a first pass, all papers with relevant images were visually inspected without the aid of anything except the occasional rotation and contrast adjustment. Following this first pass, we were granted beta access to Imagetwin (Imagetwin AI GmbH, Vienna, Austria) – online image analysis software designed to detect duplications. This allowed us to detect also duplications between publications, something that is mostly not possible for a human analyst. Consequently, we undertook a second, software-assisted, pass of all papers with relevant images.

### Classification

The issues we found in published images presenting research data were classified according to the system developed by Bik *et al*.^15^ (refer to **Figure 1**). Each publication that we screened was listed either as “clean”, containing no image issues that we could detect, or as problematic. A small number of publications where we felt that we could not make a fair assessment of the images (for example, where the images could only be obtained at very low resolutions) were listed as neither. Grading was carried out on a publication level. Where multiple duplications were found within the same paper, we associated the publication with the highest classification that we found. We also recorded the types of images where we encountered issues for the problematic papers.

Two slight modifications were made to the original classification system. We found a small number of images with issues that did not fit well in the three levels of the “Bik scale.” Consequently, we added a category labelled “other.” In this category, images often contained technical issues, where they could not have been obtained in the way outlined in the report. A typical example is a situation where two sets of western blots were allegedly imaged from the same membrane, but where this was clearly not possible. The second modification concerned duplications between reports. The classification system was originally developed for images viewed only in the context of the publication they were found in. When we found that authors had duplicated an image between publications, we upgraded this to a Type II duplication even if no repositioning had occurred. We believe this to be in the spirit of the original classification system, as this could not be considered a “simple” duplication. A lot more would have to go wrong for this to happen by accident. Moreover, when a duplication was found between reports where there were no shared authors, we listed this as a Type III duplication. Only the last published report (going by year of publication) was listed as problematic, seeing as we had no reason to believe that anything had gone wrong with the original publication. We did however exclude also the original publications from our list of “clean” studies in our bibliometric analysis.

### Reporting

All image issues were reported, over a period of two years, on the post-publication peer review platform PubPeer (www.pubpeer.com), with the exception of five papers, where the digital object identifier (DOI) and/or PubMed IDs of the papers were not compatible with PubPeer. Two papers were found already to have issues reported on PubPeer by anonymous user *Hoya camphorifolia*. Where we reported on issues with papers, anonymous users (users: *Hoya camphorifolia, Mycosphaerella arachidis, Schinia honesta*, and *Uromycladium fusisporum*) made additional discoveries in five cases. In a number of cases, anonymous user *Illex illecebrosus* provided animations demonstrating some of the overlapping images we had listed on the YouTube channel ZeebaTV. Responses submitted through PubPeer were recorded in August 2023.

### Bibliometrics

Outside of the metadata obtained in the systematic review, we recorded the authors’ national affiliation(s) for each of the 1,035 publications.

A list of 447 “clean” publications, with no remarks on the images, was constructed. A matched-size sample of 112 papers was randomly selected (using a random number generator) and contrasted in a bibliometric analysis with the papers flagged with image issues. The journals that published the 112 “clean” and 112 papers with problematic images were noted and their journal impact factor was obtained through Web of Science (Clarivate). We also obtained the number of citations to each of these items, also through Web of Science. Bibliometrics were obtained in July 2023. To usefully contrast the number of citations to papers published as much as fourteen years apart, we calculated the average number of citations per year. Papers published in journals with no impact factor listed for 2022 (n = 29) were excluded from analyses (rather than using an impact factor from a previous year, which was sometimes an option). Only one publication in our analyses had never been cited.

### Meta-analysis

A total of 132 studies (describing 171 experiments relevant for our research question) met our pre-registered criteria for being included in a systematic review. Of those studies, ten were flagged with problematic images. One study reported on two separate experiments, both contributing evidence to our primary research question. As a consequence, eleven experiments in our meta-analysis were associated with reports containing problematic images. This subset of experiments was compared to the other 160 experiments in the systematic review that were not associated with problematic images. The individual effect sizes (expressed as standardized mean differences – Hedges’ g) were pooled in a random-effects model using the DerSimonian and Laird method for estimating between-experiment heterogeneity. The subgroups were subsequently compared in a Q-test assessing the hypothesis that the effect sizes between the two cohorts differed significantly. The analyses were carried out in R Studio (RStudio Team, 2022) running R v. 4.2.1 (R Core Team, 2022) using the package “meta”^62^.

## Supporting information

Supplemental Materials (raw Data)

## Supplemental materials

For a complete list of studies included in the analysis, a breakdown of the flagged papers with links to the individual PubPeer entries, the data used in the bibliometric analysis and quality of reporting/risk of bias analyses, please refer to the supplemental materials. For additional information with respect to the methods, refer to the original publication (https://doi.org/10.1038/s41398-024-02742-0). For raw data used for the meta-analytic synthesis of effect sizes and analysis code, refer to material available through the Open Science Framework (https://osf.io/wdba5).

## Acknowledgements

We would like to thank the Danish 3R Center that supported our original work (grant number 33010-NIFA-20-743), Patrick Starke and the team at ImageTwin AI GmbH for allowing us early access to their software, and Dr. James Heathers for offering some consultation on the topic at hand.

